# Non-invasive evidence for rhythmic interactions between the human brain, spinal cord, and muscle

**DOI:** 10.1101/2024.05.01.591590

**Authors:** Meaghan E. Spedden, George C. O’Neill, Ryan C. Timms, Timothy O. West, Stephanie Mellor, Tim M. Tierney, Nicholas A. Alexander, Robert Seymour, Simon F. Farmer, Sven Bestmann, Gareth R. Barnes

## Abstract

Voluntary human movement relies on interactions between the spinal cord, brain, and sensory afferents. The integrative function of the spinal cord has proven particularly difficult to study directly and non-invasively in humans due to challenges in measuring spinal cord activity. Investigations of sensorimotor integration often rely on cortico-muscular coupling, which can capture interactions between the brain and muscle, but cannot reveal how the spinal cord mediates this communication. Here, we introduce a system for direct, non-invasive imaging of concurrent brain and cervical spinal cord activity in humans using optically-pumped magnetometers (OPMs). We used this system to study endogenous interactions between the brain, spinal cord, and muscle involved in sensorimotor control during simple maintained contraction. Participants (*n*=3) performed a hand contraction with real-time visual feedback while we recorded brain and spinal cord activity using OPMs and muscle activity using EMG. We first identify the part of the spinal cord exhibiting a peak in estimated current flow in the cervical region during contraction. We then demonstrate that rhythmic activity in the spinal cord exhibits significant coupling with both brain and muscle activity in the 5-35 Hz frequency range. These findings evidence the possibility of concurrent spatio-temporal imaging along the entire neuro-axis.

## 1. Introduction

Interactions between the spinal cord, brain, and sensory afferents shape the control of voluntary movement [1], [2], [3]. The spinal cord integrates the activity of afferents with descending motor commands to precisely fine-tune the activation of motor neurons, representing the final common pathway for transmitting neural information to the muscle [4], [5], [6]. Hence, the spinal cord serves an integrative function critical for natural movement [2], [7]. As yet, this integrative function has proven particularly difficult to study in humans, in part due to methodological challenges associated with concurrent spatio-temporal imaging of brain and spinal cord activity.

A new generation of magnetic sensors, termed optically pumped magnetometers (OPMs), offer remarkable flexibility in sensor array construction, which allows us to study the spatio-temporal dynamics of the nervous system as a whole. OPMs are small, lightweight sensors which, depending on the insulation provided, can be placed very near skin (∼10 mm or closer) [8], [9], [10], [11]. This also means that sensor placement is flexible, which makes it feasible to construct custom arrays covering the head and back to record brain and spinal cord activity simultaneously. Critically, an OPM-based system for concurrent brain (magnetoencephalography, MEG) and spinal cord imaging (magnetospinography, MSG) has the potential to give us a direct but non-invasive spatial and temporal window into spinal cord activity and how it interacts with the brain and muscle. We recently introduced such a magneto-spino-encephalography (MSEG) system by showing that we can measure evoked responses at expected latencies from the spinal cord and brain concurrently, using peripheral nerve stimulation [12].

Building on this work, we test the utility of OPMs in assessing the spatio-temporal dynamics of interactions between the brain, spinal cord, and muscle, as a basis for voluntary motor behaviour. In the sensorimotor system, a large body of research has focused on cortico-muscular coherence, which comprises a linear, within-frequency phase and amplitude coupling between brain and muscle activity [13]. Cortico-muscular coherence is present during low-level tonic contraction of distal muscles and exhibits a characteristic peak in the beta band (15-30 Hz) [14], [15], [16], [17]. It can also be detected as a common drive to individual motor neurones, both within and between muscles [15], [16]. This coherence thus reflects the transmission of rhythmic activity between the brain and muscle, through the spinal cord, likely to involve both ascending and descending pathways [18].

Consequently, our understanding of interactions between the brain, spinal cord, and muscle during movement has been largely based on inferring spinal cord activity indirectly from muscle activity. Based on previous work [14], [16] we can infer that the rhythmic activity reflected in cortico-muscular coherence (and other functional connectivity measures) is relayed through neurons in the spinal cord, and that these neurons can affect and be affected by rhythmic neural activity. Direct evidence for this comes from non-human primate studies reporting coherence between spinal cord local field potentials and muscle activity during a precision grip task [19], [20]. The ability to measure such interactions directly, and non-invasively remains a goal for human neuroscience.

Here we address this need through direct, concurrent, and non-invasive measurements of cortical and spinal activity using a custom-built array of OPMs. We quantify rhythmic coupling between brain and spinal cord activity (measured with OPMs), and muscle activity (measured with electromyography, EMG) during a simple, tonic hand muscle contraction using metrics of interaction on signal phase or signal amplitude.

## 2. Methods

### 2.1 Participants

Three healthy, right-handed male participants took part in this series of experiments (participant A, aged 45; participant B, aged 55; participant C, aged 28) after giving written, informed consent. The study procedures were approved by the University College London Research Ethics Committee and experiments were conducted in accordance with the Declaration of Helsinki. Informed consent has been obtained from all study participants for the publication of images in an online open access publication.

### 2.2 Summary

Participants performed a tonic thumb contraction task where they were asked to maintain a constant low-level contraction as precisely as possible using real-time visual feedback of their muscle activity. While they performed this task, we used OPMs to record activity from the brain and spinal cord, and EMG to record muscle activity, with the aim of studying functional interactions between the three regions.

### 2.3 OPM recordings

For this study, we developed an OPM-based system to perform concurrent recordings of brain and cervical spinal cord activity while participants lay supine.

#### 2.3.1 Head and neck casts

Second and third generation QuSpin (Louisville, CO) manufactured OPM sensors (dual-axis and triaxial sensors, respectively) were positioned in custom-built, 3-D printed (Chalk Studios, London, UK) rigid nylon casts for the head and neck, with the aim of recording brain and spinal cord activity. The head casts were constructed based on participant A and B’s anatomical MRI scans. Participant C did not have a custom head cast, but had a similar head size to B, and so was able to use participant B’s head cast. A generic neck cast was used for all three participants. The neck cast was designed based on the shape of the Siemens 64-channel Head and Neck MRI coil (Siemens Healthineers, Erlangen, Germany) to fit a range of head and neck sizes and shapes. On the lower-surface of the cast, we added 42 slots covering the upper back, neck, and bottom of the head. We populated the neck cast with sensors as fully as possible (i.e., depending on sensor availability), then placed on average 13 sensors around the right and left sensorimotor cortex region in the head casts (Supplementary Table 1).

#### 2.3.2 OPM acquisition

Experiments were conducted in a magnetically shielded room (MSR) (438 x 338 x 218 cm; Magnetic Shields Ltd, Staplehurst, UK). The inner layer of mu metal lining the room was degaussed using a low-frequency decaying sinusoid driven through cables within the walls prior to the start of the experiment. The OPM sensors were then nulled using on-board nulling coils and calibrated. OPM data were acquired using a National Instruments acquisition system and a LABVIEW program with a sampling frequency of 6000 Hz and 16-bit resolution. An antialiasing 500 Hz low-pass filter (60th order FIR filter combined with a Kaiser window) was applied before data were down-sampled offline to 2 kHz.

### 2.4 EMG recordings

EMG was recorded from the right and left abductor pollicis brevis muscle (APB) using pairs of surface electrodes (Ambu Ltd, Ballerup, Denmark). The APB is innervated by the median nerve, which enters the spinal cord around C6-T1 [21]. The reference electrode was fastened around the right wrist. In order to prevent EMG data collection from interfering with OPM acquisition, we passed EMG cables through waveguides so that EMG signals were amplified, filtered, and digitized outside the MSR (Amplification: x 1000; filtering 3 to 1000 Hz; digitization at 2000 Hz; D-360 Amplifier, D-400 Mains noise eliminator, and 1401 data acquisition unit, Cambridge Electronic Design, UK). EMG data were recorded using Spike2 software (v10.05). EMG and OPM data were synchronized based on a TTL pulse sent from the 1401 data acquisition unit.

### 2.5 Task

During recordings, participants lay on their back on a cushioned, plastic bed in the MSR wearing a head cast and with their head resting in the neck cast (Figure 1).

**Figure 1.**
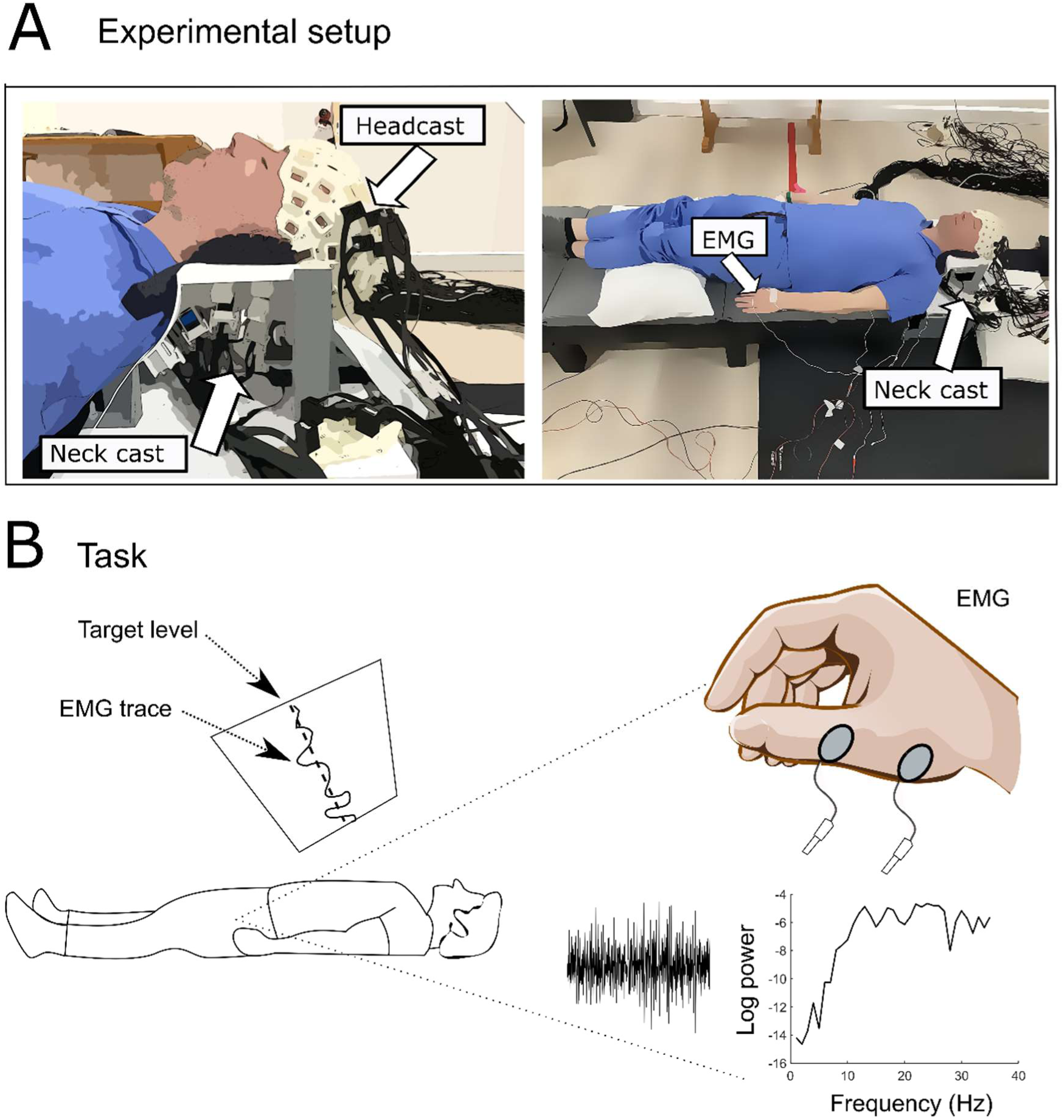
Experimental setup. A. Participants lay supine on a plastic bed in the magnetically shielded room wearing a head cast (holding the cortical OPM sensors) and with their head and neck resting in a neck cast (holding the spinal cord sensors). The casts were populated with dual– and triaxial OPMs (on average 29 sensors in neck cast and 13 in head cast; see Supplementary Table 1). EMG was recorded from the right and left short thumb abductors. B. Participants were shown a target contraction line corresponding to ∼10 % of their maximal voluntary contraction EMG projected onto the ceiling above them. The goal of the task was to track this target line with their real-time rectified, smoothed EMG trace as precisely as possible.

We selected a weak isotonic contraction of a distal muscle because it elicits the well-described beta band coherence between the brain and muscle [14], [22], [23], providing a helpful point of departure for further investigating functional connectivity between the brain, spinal cord, and muscle.

We projected a target as a horizontal line onto the ceiling above the participants, along with a trace of their real-time rectified, smoothed APB EMG signal. Participants were asked to follow this target level with the real-time EMG trace as precisely as possible for two minutes at a time using a rubber ball as resistance for the contraction. The target level was determined based on 10 % of the rectified, smoothed APB EMG trace during a maximal voluntary contraction. Before beginning the recordings, we tested that the target level was comfortable for the participant to maintain for longer periods, and based on feedback we lowered the level slightly for participant C. Four 2-minute runs were recorded for the right thumb, interspersed with two minutes of rest to avoid fatigue. The task was subsequently repeated for the left thumb. For participant A only three runs were recorded for the left contraction. Total experiment duration was approximately 40 minutes. Task performance was quantified as the root mean square deviation of the rectified smoothed EMG trace from the target level (Supplementary Table 2).

### 2.6 Data analysis

The main outcomes of the analysis comprise (1) source reconstructed spinal cord data, used to define a region of interest based on the point of maximal estimated current flow for each participant; and (2) metrics of functional connectivity between brain and muscle activity; muscle and spinal cord activity; and brain and spinal cord activity.

All data analysis was performed in MATLAB R2021b (Mathworks Inc., Natick, MA, USA) using the development version of Statistical Parametric Mapping (SPM; https://github.com/spm) [24] and Fieldtrip [25] toolboxes.

#### 2.6.1 Pre-processing of OPM and EMG data

In the following, we use the terms ‘sensor’ to describe one dual or triaxial OPM sensor and ‘channel’ to describe the signal from one sensor axis. OPM data were read into SPM and power spectra and time series were inspected for evidence of bad channels (deviating from the median power spectrum; saturating; or showing consistent and large amplitude deflections over time), which were removed from the data set. OPM data were high pass filtered at 5 Hz, then low pass filtered at 45 Hz, using 5^th^ order Butterworth filters, then 3^rd^ order band stop filters from 48-52 and 98-102 Hz were applied. EMG data were high pass filtered at 10 Hz using a 5^th^ order Butterworth filter. Using three reference sensors positioned on the support structure for the neck cast, i.e., away from the participant (approximately 15 cm cranial to and 20 cm lateral to the participant’s inion), we then performed synthetic gradiometry [26], regressing channels from the reference sensors out of brain and spinal cord channels. The reference sensors capture environmental interference, but not neural signals; consequently, the synthetic gradiometry subtracts this interference from the signals of interest.

After filtering the OPM data, large heartbeat artefacts were apparent on most channels. We removed these artefacts using signal space projection (SSP) [27]. First, we used singular value decomposition (SVD) of the covariance matrix for the spinal cord channels to obtain a component reflecting the heart beat artefact time course. Heart components were selected among the first four components based on visual inspection of their time series. The selected component time course was used to create an artefact template comprising averaged OPM data time-locked to the main peak of the artefact (assumed the ‘R’ in the QRS complex). Data were then segmented into non-overlapping epochs with one second duration and trials were classified as outliers if the mean variance of the trial over channels exceeded a threshold of 3 standard deviations above the median mean trial variance. Outlier trials were removed, and remaining trials were merged for right and left contractions, respectively. Finally, data were projected into a subspace orthogonal to the heartbeat artefact direction. The lead fields were pre-multiplied by the SSP projector [27] to account for any distortions induced by the SSP projector in the spinal cord signal.

#### 2.6.2 Estimates of current flow in the spinal cord

We conducted source analysis on the OPM data recorded from the sensors in the neck cast to estimate current flow in the spinal cord.

We performed optical scans (SHINING 3D Tech. Co., Ltd., Hangzhou, China) of each participant’s head, neck, and upper back (without the head or neck casts) while they were seated. This provided a head and neck shape into which a ∼20 mm diameter cylinder was placed at the approximate depth of the cervical spinal cord. We approximated current flow within this cylinder as arising from dipolar sources spaced at 10mm intervals (the number of source points totalled approximately 145). Dipolar sources were oriented in the dorsal-ventral direction. During this type of task requiring integration of proprioceptive feedback with motor output, there is likely current flow along the cord, in the medial-lateral direction, and in the dorsal-ventral direction [28], but we chose this orientation because it presumably arises from spinal cord grey matter, which we consider instrumental for sensori-motor integration in the spinal cord.

The optical scan was co-registered to the sensor positions by manually registering the points in the optical scan to the surface of the neck cast, based on photos and a third optical scan of participants lying down, wearing the casts during the experiment (see Figure 2 for co-registered output). For participant C it was not possible to obtain a scan lying down and wearing the casts, so photos of the X Y and Z axes were used to guide the co-registration using Blender software (v3.3.1).

**Figure 2:**
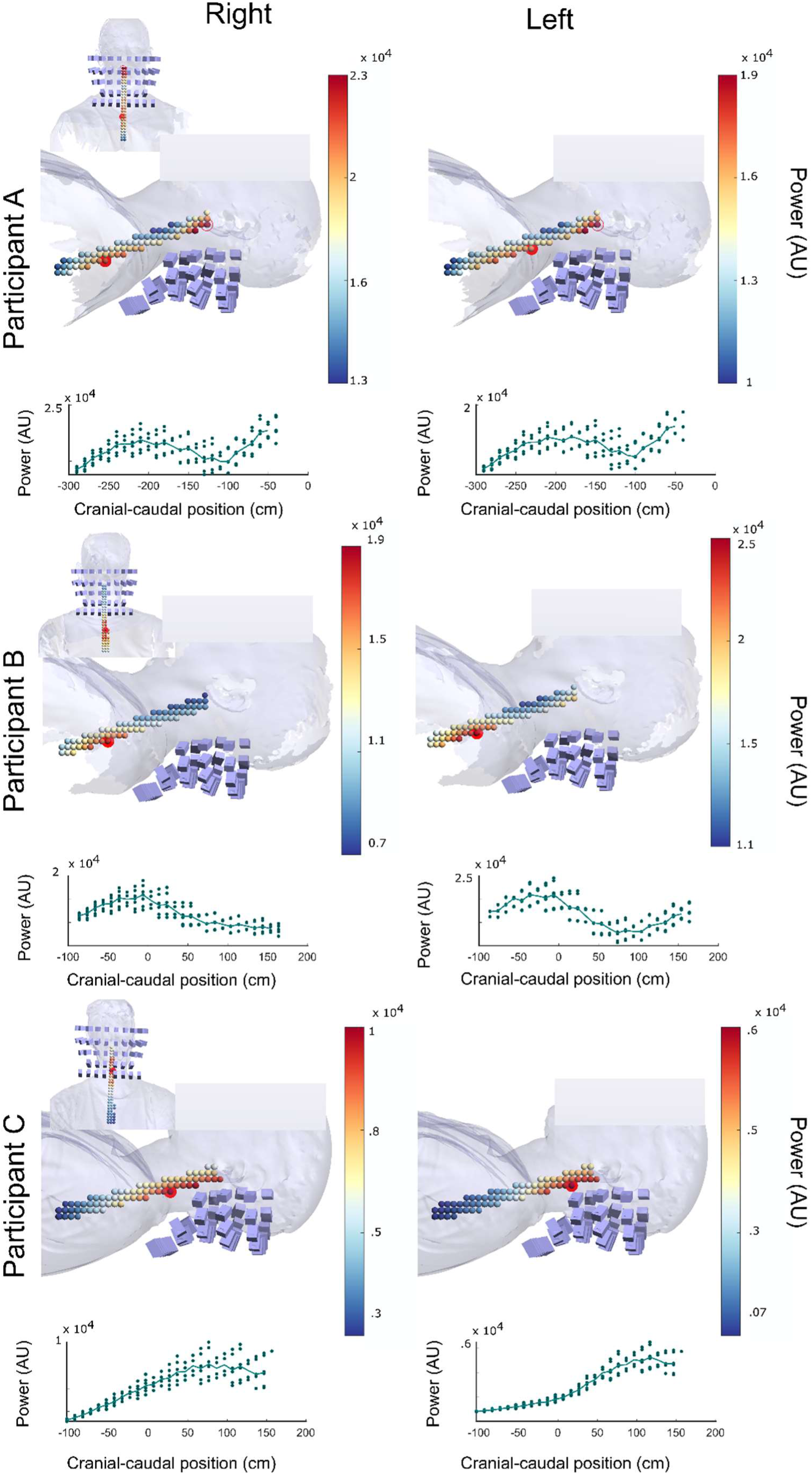
Source reconstructed spinal cord activity during right and left thumb contraction. Participant head and neck shapes constructed using optical scans are shown in light blue. Purple rectangles display sensor positions. The source space was a cylindrical grid of dipoles (small spheres). Sphere colour reflects the estimated power at each source point. The point of maximal power in the cervical region is highlighted with a filled red dot. For participant B this was different from the global maximum (highlighted with a red circle). Reconstructed source power is also illustrated as a function of cranial-caudal position in source space (bottom thumbnails).

This procedure gave us the approximate sensor positions and orientations in the same coordinate space as the source grid. We used an infinite homogenous medium as our volume conductor because it has been successfully used previously to reconstruct spinal cord activity recorded with cryogenic sensors [29], [30]. Lead fields were calculated based on the source grid, sensor positions and orientations, and the infinite volume conductor in Fieldtrip.

This forward model was used to perform a volumetric Bayesian minimum norm inversion in SPM (‘IID’) [24], [31] to estimate current flow in the spinal cord. We used frequency domain spinal cord data from the tonic contraction as input to the source reconstruction. We chose a frequency range of 5-35 Hz to cover alpha/mu and beta frequencies, as these bands show strong movement-related modulations in the sensorimotor system [13]. Fourier coefficients were calculated using the multitaper method using discrete prolate spheroidal sequences (DPSS; *n*=5 tapers) [32]. Specifically, input to the source analysis were Fourier coefficients for *Nc* channels, *m* features (= number of frequencies x 2; real and imaginary parts) and *Nt* (=number of trials): data matrix *X* ϵ ℝ^Nc^ ^x^ ^m^ ^x^ ^Nt^.

We performed the source reconstruction for right and left thumb contraction for each participant to get the maximum a posteriori projector to map between spinal sources and sensor space, then used this mapping to calculate trial-wise complex Fourier coefficients in source space for our frequencies of interest. Source activity is summarized as the power (current density squared) at each source point. We defined our region of interest for subsequent functional connectivity analyses to be the location of peak power (in the 5-35Hz band) in the cervical region.

#### 2.6.3 Brain signal extraction

Because we used relatively few sensors to record brain activity, we summarized activity as the first principal component (i.e. dominant spatial mixture) of brain channel covariance for the real part of the cross-spectrum between the brain and EMG. Results were similar when using the imaginary part.

Note that using the mixture of brain channels that best explains the brain-EMG cross spectrum biases our analyses towards finding cortico-muscular functional connectivity. However, our focus in this study is on detecting functional connectivity with the spinal cord, and isolating a component of the brain signal best explaining the EMG gives a robust foundation for this analysis.

#### 2.6.4 Functional connectivity analysis

We used the canonical variates analysis (CVA) framework implemented in SPM to quantify functional connectivity between 1) brain and muscle activity; 2) spinal cord and muscle activity; and 3) brain and spinal cord activity. The analysis comprised three stages, each evaluating a different functional connectivity feature: within-frequency coupling at constant phase; within-frequency amplitude envelope coupling; and cross-frequency amplitude envelope coupling.

We first give a broad overview of CVA before detailing each type of functional connectivity analysis performed using this framework. CVA is a multivariate method that can capture associations between two sets of data [33], [34], [35], [36].

Given two multivariate datasets *X* ϵ ℝ^Nt×m^ and *Y* ϵ ℝ^Nt×n^ with *m* and *n* features (or columns) respectively over *Nt* trials (or rows), CVA finds linear combinations of each of the two data sets that maximize the correlation between them. These linear combinations (or canonical vectors) are *V* ϵ ℝ^m×q^ and *W* ϵ ℝ^n×q^ where *q=min(m,n)*, such that the best linear prediction of the multivariate mixture *YW* is *XV* (the columns of these matrices are often referred to as canonical variates).

Or alternatively, the prediction of *Y* based on *X* is 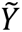 where

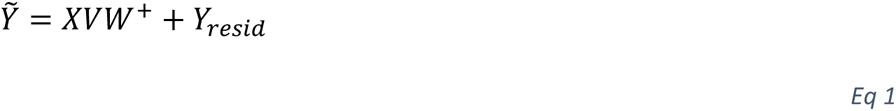

Where *W^+^*signifies the Moore-Penrose pseudo-inverse of *W* and the residual (unpredicted) portion of *Y* is *Y*_resid_ ϵ ℝ^Nt×q^.

This framework can also account for a null space (or set of confounds) *X0* which is treated as a regressor of no interest. All data were mean-centred before entering them into the CVA.

Statistically, CVA tests the hypothesis that there exist linear mappings from Y to X providing an *R^2^*test statistic and *p*-value for each of *q* dimensions or canonical modes of correlation.

With degrees of freedom for ith mode *(m-i+1)*(n-i+1)*, where *m* and *n* are the columns (or features) in *X* and *Y* respectively.

We then used these tests to separately analyse coupling between neural signals: within-frequency at constant phase; within-frequency amplitude envelope coupling; and cross-frequency amplitude envelope coupling.

All functional connectivity analyses were performed on frequency domain data, i.e., trial-wise Fourier coefficients for our frequency range of interest (5-35 Hz) calculated using the DPSS multitaper method.

##### 2.6.4.1 Within-frequency phase locking

We began our functional connectivity analysis by considering the consistency of (real and imaginary) phase coupling across *Nt* (=number of trials x number of tapers). For each frequency, input to the CVA were data *Y* ϵ ℝ^Nt×2^ and *X* ϵ ℝ^Nt×2^. The two columns of the data contained the real and imaginary parts of the Fourier coefficients at a single frequency. For analysis of brain-muscle functional connectivity, *Y* was the optimal brain signal and *X* was EMG; for muscle-spinal cord functional connectivity, *Y* was EMG and *X* was spinal cord activity at the point of maximal reconstructed current flow; and for brain-spinal cord functional connectivity, *Y* was the optimal brain signal and *X* was spinal cord activity at the point of maximal reconstructed current flow. This multivariate test was run at each frequency.

This metric is based on assessing whether there exist linear mappings between the real and imaginary parts of X and those of Y. Importantly, the classical multivariate framework allows us to include a comprehensive null space, returns any linear mapping between X and Y and, in addition to a metric of association (χ^2^, R^2^), a significance value. We used a null space *X0* ϵ ℝ^Nt×(2C)^ containing *C* (real and imaginary) principal components of the reference channel Fourier coefficients that explained 95 % variance of the reference channel data to control for any shared variance not removed by the synthetic gradiometry. The cost of the multivariate approach is that there are effectively 4 parameters estimated from the data and degrees of freedom removed by the null-space (i.e. degrees of freedom in the residuals = *Nt-n-m-2C*) [33]. In this case, for *Nt* >> *n*+*m*+*2C* (*n*=2, *m*=2, C < *Nc*) with number of reference channels *Nc*, the reduction in the effective degrees of freedom does not present an issue and there is little danger of over-fitting. The advantage of the extra association parameters estimated is that the method is robust to interfering signals which may impact conventional coherence metrics. For example, consider a signal in which the real part of *X* predicts the imaginary part of *Y*; but the real part of *Y* is a pure noise term. The multivariate method has the flexibility to identify this mapping, yet *X* and *Y* would have low coherence due to the noise term (perturbing the phase of *Y*). (see Supplementary Figure 6 and associated text for an example).

Results were quantified as *R^2^* with a corresponding *p*-value of the χ^2^ statistic testing that there exists (at least one) linear mapping from *Y* to *X* each frequency. Here we only report results for the first canonical mode of correlation, i.e., *q*=1. We corrected for multiple comparisons by using a False Discovery Rate (FDR) correction at 0.05 [37].

##### 2.6.4.2 Within-frequency amplitude envelope coupling

We then turned to evaluate within-frequency amplitude envelope coupling. Here, we used the residuals from our phase-based analysis (*Y_resid_*) with the aim of ensuring that any amplitude changes in *Y* predicted from *X* due to a constant phase relationship does not influence the estimate of envelope-based coupling.

We looked to see if the envelopes (i.e. purely the trial by trial signal amplitude) of *X* and *Y* were related at any frequency. Note that this is simply a linear regression, but we keep to the CVA notation for consistency.

We took the absolute value of *Y_resid_* and *X* to quantify coupling between signal amplitude envelopes. Specifically, at each frequency, input to the CVA were *Y* ϵ ℝ^Nt×l^ and *X* ϵ ℝ^Nt×l^. In order to avoid common environmental noise factors, we used the mean corrected envelope of the sparse reference set *X0* (above) as a confound *X0_env_* ϵ ℝ^Nt×C^. Results were quantified as *R^2^* with a corresponding *p*-value at each frequency. We corrected for multiple comparisons by using an FDR correction at *p* < 0.05. The within-frequency envelope prediction of *Y_resid_* from *X* was removed to give envelope residuals *Y_resid_env_*.

##### 2.6.4.3 Cross-frequency amplitude envelope coupling

In the final stage of the functional connectivity analysis, we used the residuals from the envelope analysis *Y_resid_env_* ϵ ℝ^Nt×f^ and data matrix *X* ϵ ℝ^Nt×f^ as input to the CVA to investigate cross-frequency envelope coupling. In this analysis, CVA finds the mixture of envelopes (across frequencies) for each data matrix that are most strongly correlated. We report the results of this analysis as the *p*-value for the first mode of canonical correlation. Results are also visualized across frequencies in Supplementary Figures 4-5 as the transformed canonical vectors (*V,W*) [38] determining the linear-mixing of frequency envelopes that predicts *Y* from *X*.

##### 2.6.4.4 Relationship between functional connectivity and task precision

Finally, we explored whether task precision played a role in the preceding functional connectivity estimates. For each of the above analyses we added an extra column to *X*. The column contained a metric of task performance on each trial. Task performance was quantified as the root mean square (RMS) error for the produced rectified, smoothed EMG relative to the target level for periods of 1 second. This reflects how precisely participants were able to maintain the target EMG level. Adding an extra predictor inevitably increases the variance explained; however, there is a principled way of testing whether the increase in χ^2^ is significant [33]. Using the ratio of χ^2^ values for the first canonical mode of correlation with and without the predictor, normalized by their degrees of freedom, we can calculate an F-statistic and corresponding *p*-value [39]. *P*-values were corrected using FDR for within-frequency phase and amplitude envelope analyses as tests were performed at each frequency of interest.

## 3. Results

We used OPMs to record brain and spinal cord activity during tonic thumb contraction in three participants. First, we present the results of our source imaging of spinal cord activity. Next, we quantify and characterize functional connectivity between (1) the brain and muscle; (2) the muscle and spinal cord; and (3) the brain and spinal cord.

### 3.1 Reconstructed spinal cord current flow shows a peak in the cervical region

We first sought to identify a region of interest for our functional connectivity analysis by reconstructing source-level spinal cord activity and examining its spatial distribution. The APB muscle is innervated by the median nerve, which enters the spinal cord around C6-T1; consequently, we expected to detect a peak in activity roughly in this region during thumb contraction.

Figure 2 shows the source reconstructed spinal cord activity for each of the three participants during right and left contraction. All participants showed a peak in power around the cervical region of the spinal cord. The spatial maps of spinal cord activity during right and left thumb contraction exhibited clear similarities, although based on independent data.

We used the point of maximal estimated source power in the cervical region as a region of interest for further functional connectivity analyses. For participants B and C this was also the global maximum, but for A we chose the local maximum of the more caudal peak to ensure we were capturing spinal cord activity. Note that the choice of selection criterium for the region of interest (power) is independent of the choice of the interaction statistic (functional connectivity).

### 3.2 The brain, spinal cord, and muscle interact during tonic contraction

Having established our spinal cord region of interest, we set out to quantify how the brain, spinal cord, and muscle interact during tonic contraction. We used CVA to quantify functional connectivity between (1) the brain and muscle; (2) the spinal cord and muscle; and (3) the brain and spinal cord. We quantified three types of coupling: within-frequency phase locking, within-frequency amplitude envelope coupling, and cross-frequency amplitude envelope coupling. For simplicity we focus on results from right thumb contraction and show left contraction results in Supplementary Figures 1-3. Results were largely similar for right and left contraction, although based on independent data, and we note any deviations in the text. We begin the analysis by looking at and replicating previously accessible measures of brain-muscle interaction; we then focus on the unique contribution of this paper which is to examine cord to muscle and cord to brain interactions.

#### 3.2.1 Interactions between the brain and muscle

We started by asking whether brain and muscle activity exhibited within-frequency phase locking in our frequency range of interest (5-35 Hz). These results are presented in Figure 3A-C. Note that we visualize coupling values from 1-35 Hz to show all of the low frequency data features (i.e. not showing these features results in a hard edge at 5Hz), but we consider our statistical tests for the 5-35 Hz range, encompassing alpha and beta bands known to exhibit movement-related modulations.

**Figure 3:**
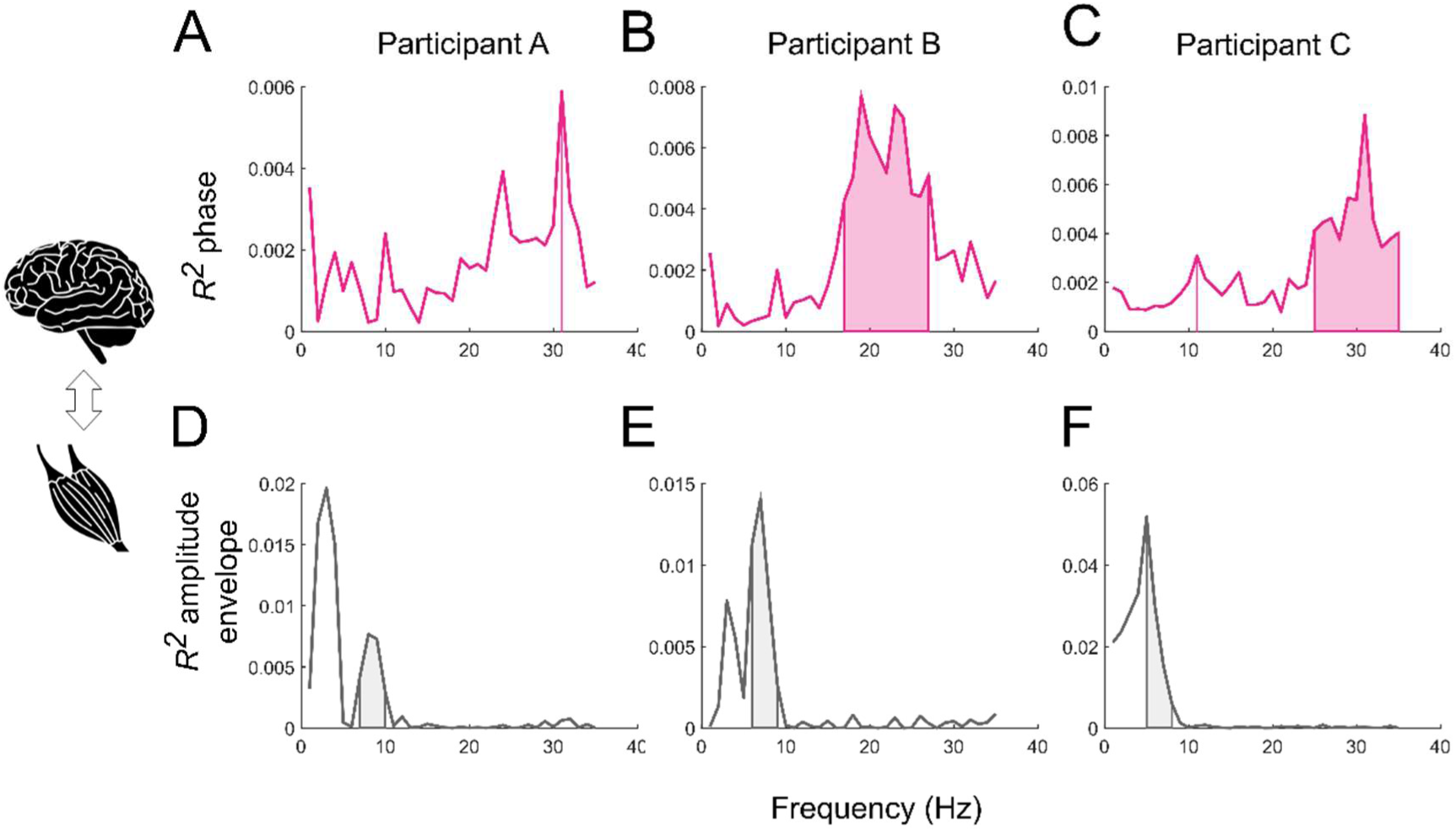
Functional connectivity between brain and muscle activity during right thumb contraction. *R^2^* for within-frequency phase locking (**A-C**). Vertical lines and shaded areas indicate FDR corrected *p*<0.05. Note that participants exhibit peaks in the beta band suggesting the synchronization of rhythmic activity at these frequencies. **D-F**: *R^2^* for within-frequency amplitude envelope coupling. These results reflect coordinated fluctuations of amplitude envelopes at frequencies < 10 Hz. Y-axes are scaled differently to clearly show spectral features.

All three participants showed significant peaks in canonical correlation between brain and muscle activity at beta frequencies, indicating significant cortico-muscular phase coupling around 20-30 Hz (corrected *p* < 0.05). The beta frequencies for the peaks differed between participants within this band, which is typical for cortico-muscular phase-based coupling measures [40]. This suggests consistent phase relationships across trials. Note however that the coupling did not reach statistical significance for the left/non-dominant hand (Supplementary Figure 1). These results broadly support that rhythmic activity in the brain and muscle synchronize at beta frequencies during tonic contraction, at least for the dominant hand.

We then asked whether the brain and muscle exhibited amplitude envelope coupling in our frequency band of interest. For contraction of the right thumb, we found significant coupling at frequencies around 5-10 Hz across participants (Figure 3D-F; corrected *p <* 0.05).

Finally, we used CVA to evaluate cross-frequency functional connectivity. We found evidence for significant cross-frequency amplitude envelope coupling in all three participants (all *p* < 0.001). The frequencies contributing most prominently and most consistently to this coupling were < 15 Hz and are illustrated as the transformed canonical vectors [38] as a function of frequency in Supplementary Figures 4-5. Generally, it did not appear to be the case that specific, remote frequencies tended to be coupled but rather that the analysis detected coupling between close, but not identical, frequencies, with the largest peaks below 10 Hz.

Taken together, and consistent with previous work [14], [22], our results suggest that rhythmic activity in the brain and muscle are significantly coupled during tonic contraction. The different modes of coupling exhibited different spectral signatures: phase coupling was evident at beta frequencies and amplitude envelope coupling (within– and cross-frequency) was most prominent around alpha frequencies.

#### 3.2.2 Interactions between the spinal cord and muscle

Next, we sought to address how the spinal cord interacted with the muscle during the tonic contraction. We started by considering phase relationships between the signals. Figure 4A-C shows the *R^2^* values for the coupling between reconstructed spinal cord activity at the region of interest and EMG activity. This phase coupling was not significant at any frequency for any of the three participants. Notably, the beta frequency phase locking between brain and muscle (reported in section 3.2.1) was absent, a feature we will return to in the discussion.

**Figure 4:**
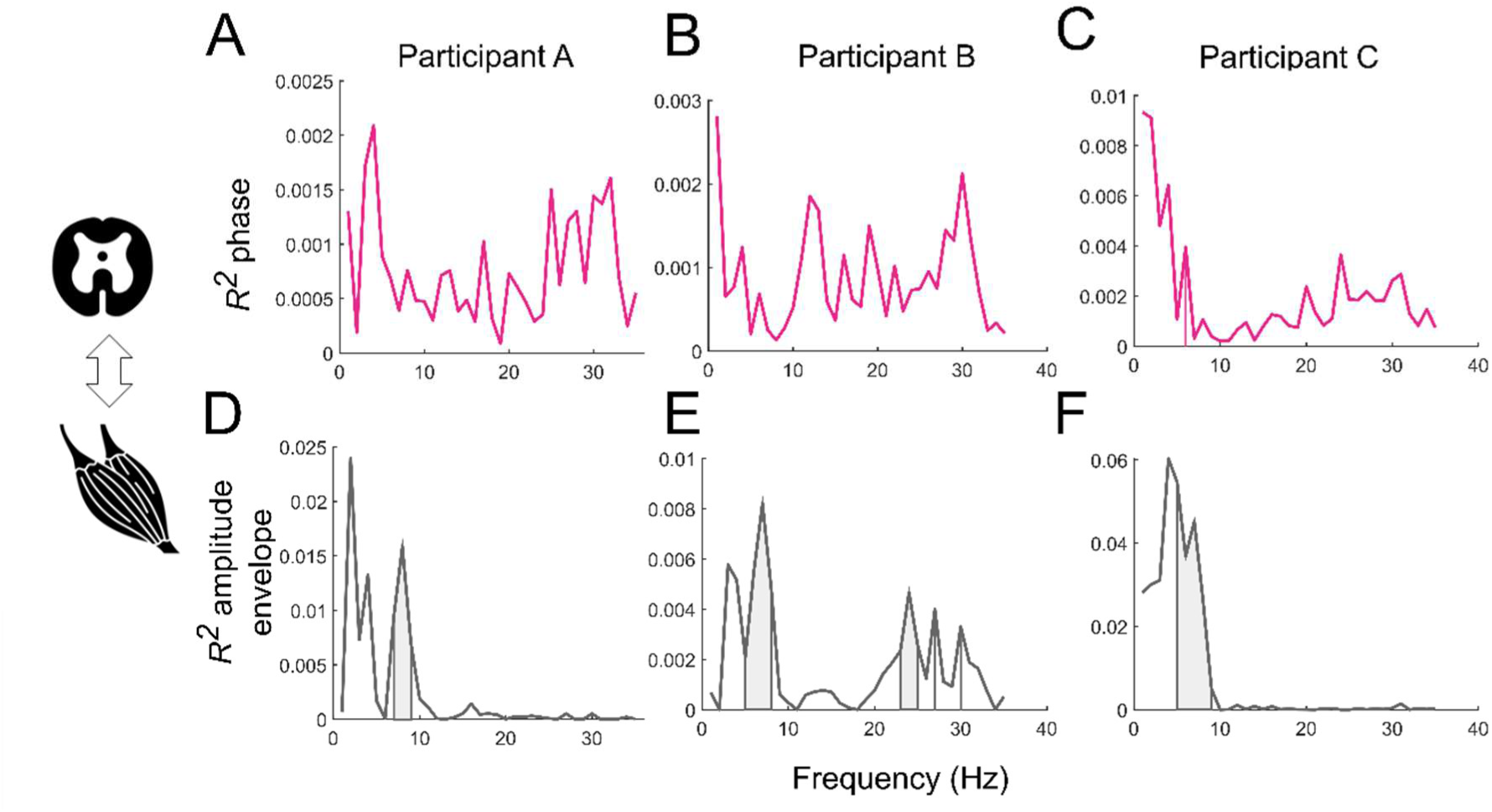
Functional connectivity between spinal cord and muscle activity during right thumb contraction. *R^2^* for within-frequency phase locking (**A-C**). Vertical lines and shaded areas indicate FDR corrected *p*<0.05. Note that the phase coupling does not reach statistical significance, suggesting that rhythmic activity between the spinal cord and muscle does not synchronize at these frequencies. **D-F**: *R^2^* for within-frequency amplitude envelope coupling. These results reflect coupled fluctuations of amplitude envelopes, most consistently at frequencies < 10 Hz. Y-axes are scaled differently to clearly show spectral features.

We then turned to evaluate within-frequency amplitude envelope coupling (Figure 4D-F). We found that source reconstructed activity at our spinal cord region of interest was significantly coupled with EMG activity at frequencies < 10 Hz (corrected *p* < 0.05), similar to what we observed for the brain-muscle amplitude envelope coupling. Participant B also showed coupling in the beta band, but this feature was not evident in the other participants’ spectra. Together, this suggests that rhythmic amplitude fluctuations in the spinal cord and muscle are coupled during the task at 5-10 Hz.

In the final stage of the functional connectivity analysis, we considered interactions across frequencies and found evidence favouring the presence of cross-frequency amplitude envelope coupling between cord and muscle in all three participants (all *p* < 0.001). The frequencies contributing to this coupling were more broadband than for the brain-muscle cross frequency coupling and are visualized in the Supplementary Material as canonical vectors (Supplementary Figures 4-5). Generally, the spectra suggested that the most prominent and consistent contributions were again from frequencies below 15 Hz.

In sum, we found evidence in this analysis for both within– and cross-frequency amplitude envelope coupling between the spinal cord and muscle predominantly at frequencies < 15 Hz, mirroring that seen when analysing coupling between brain and muscle. In contrast to results found when looking at brain-muscle interaction, spino-muscular phase locking was negligible.

#### 3.2.3 Interactions between the brain and spinal cord

Having established patterns of brain-muscle and spinal cord-muscle functional connectivity, we next extended our investigation to interactions between the brain and spinal cord.

As before, we started by quantifying within-frequency phase locking. Figure 5A-C shows that a relatively broadband association was present for participants A and C (corrected *p* < 0.05), while participant B showed multiple significant peaks throughout the frequency range rather than broadband coupling. Note however that the broadband phase-locking pattern was apparent in all three participants during left contraction, Supplementary Figure 3).

**Figure 5:**
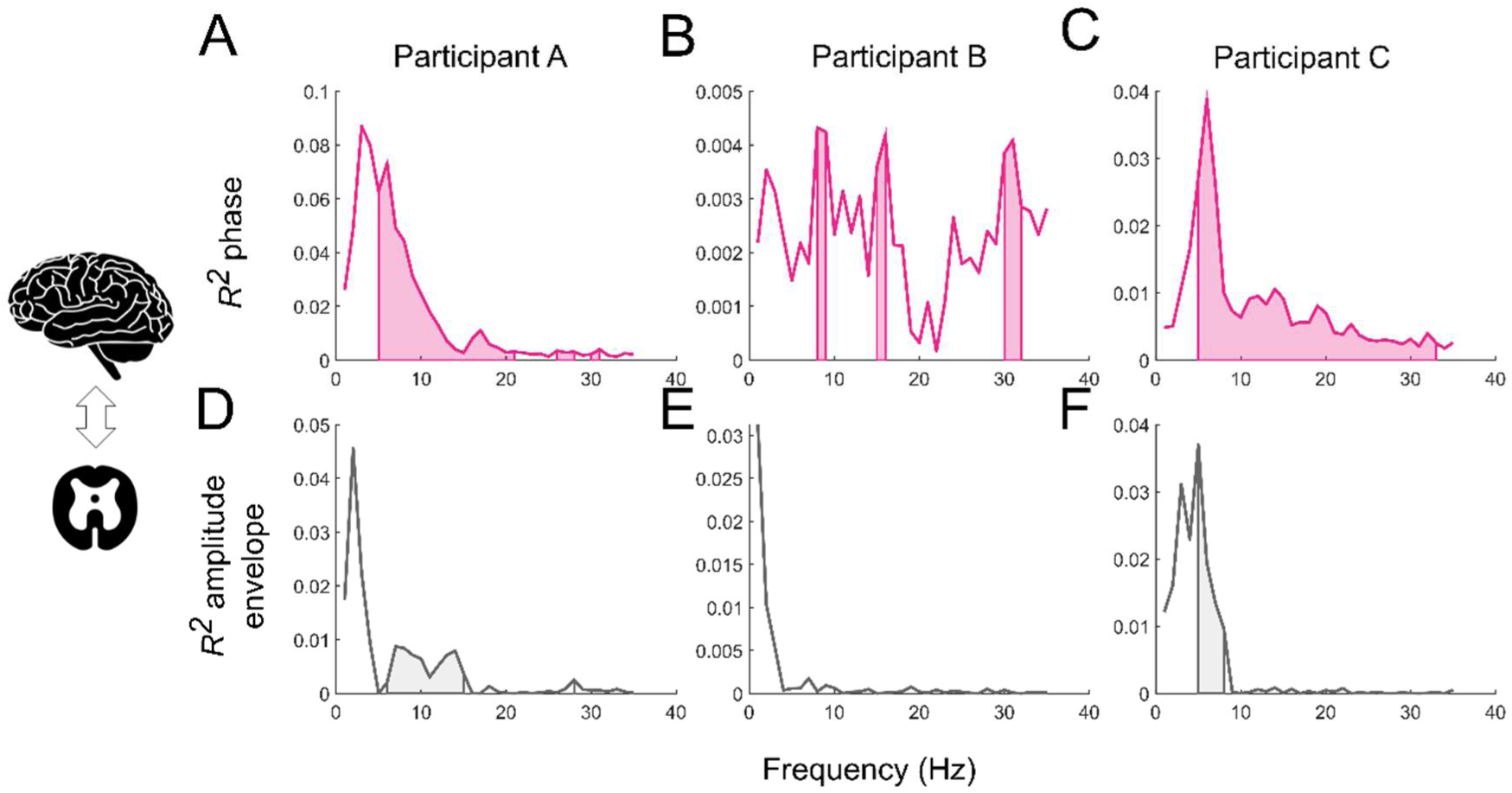
Functional connectivity between spinal cord and brain activity during right thumb contraction. *R^2^* for within-frequency phase locking (**A-C**). Vertical lines and shaded areas indicate FDR corrected *p*<0.05. Note that the synchronization occurs in both alpha and beta frequency ranges. **D-F**: *R^2^* for within-frequency amplitude envelope coupling. These results reflect coordinated fluctuations of amplitude envelopes, primarily at frequencies < 15 Hz. Y-axes are scaled differently to clearly show spectral features.

Turning to within-frequency amplitude envelope coupling (Figure 5D-F), participants A and C showed coupling at frequencies between 5 and 15 Hz (corrected *p* < 0.05), indicating that amplitude envelopes in the spinal cord and brain co-vary in this band. The lower frequency peak in participant B’s spectrum did not reach the significance level. For left contraction however, all three participants showed significant amplitude envelope coupling between 5 and 15 Hz (Supplementary Figure 3).

We then asked whether brain and spinal cord activity envelopes interact across frequencies. For the right contraction, we found significant cross-frequency coupling for brain and spinal cord amplitude envelopes for two out of three participants (A and C: *p* < 0.001; B: *p* = 0.124). Note that for left contraction, this metric was significant for all participants (Supplementary Figure 5). The most prominent frequencies contributing to this coupling were, as was the case for our previous cross-frequency envelope coupling analyses, < 15 Hz (Supplementary Figures 4-5).

Together, these results indicate that rhythmic brain, spinal cord, and muscle activity demonstrate heterogeneous functional coupling patterns in terms of phase or amplitude coupling and spectral signatures.

### 3.3 Influence of task performance on functional connectivity

Finally, having addressed functional connectivity between the brain, spinal cord, and muscle, we investigated whether task performance, quantified as RMS error of the contraction EMG relative to the target level, significantly influenced our coupling metrics.

Adding precision as a predictor to the CVA analyses did not significantly change within-frequency phase, within-frequency amplitude envelope, or cross-frequency amplitude envelope coupling for any signal pair in any participant (all corrected *p* < 0.05). Interestingly, however, we did observe multiple effects on amplitude-envelope coupling at beta frequencies that did not survive our correction for multiple comparisons (i.e., uncorrected *p* < 0.05). These uncorrected effects were most consistent across participants and hands for brain-muscle and spinal cord-muscle couplings, suggesting that these relationships may merit future exploration.

## 4. Discussion

Here, we introduce an OPM-based system for direct spatio-temporal imaging of concurrent brain and cervical spinal cord activity in humans. We show that we can non-invasively image spinal cord current flow during voluntary hand contraction, and that this rhythmic spinal cord activity co-varies with brain and muscle activity. We observed that phase and amplitude coupling exhibited distinct spectral signatures and that coupling modes differed between nodes. Together, this may hint at different underlying transfer properties of each node and signify different modes of large-scale neural transmission. We begin by discussing the source reconstruction of spinal cord activity, then proceed to discuss functional connectivity results for brain, spinal cord, and muscle interactions.

### 4.1 OPMs can be used to non-invasively image spinal cord activity

As a starting point for our functional connectivity analyses, we used source imaging to reconstruct spinal cord current flow. We found distributions of spinal cord current flow which demonstrated peaks around the cervical region of the spinal cord. This is consistent with the knowledge that our muscle of interest, the short thumb abductor, is innervated by the median nerve, which originates from the brachial plexus around C6-T1 [21]. We focused solely on where activity was localized along the cranial-caudal axis, i.e., up and down the spinal cord, but did not consider laterality. Future MSG work will hopefully be able to distinguish lateralized activity in the spinal cord, but in this early iteration, we considered similar results for right and left contraction as useful within-participant replications.

These findings are also broadly consistent with work using super-conducting magnetic field sensors to reconstruct spinal cord activity resulting from peripheral nerve stimulation [28], [30], [41]. These authors have shown that they are able to distinguish three different types of spinal cord current flow generated by peripheral nerve stimulation. They show an inward current reflecting entry to the dorsal root, ascending activity, and a posterior to anterior current presumably arising from spinal cord grey matter [28]. These results are informative in our context because they support that it is possible to detect fields arising from both travelling and static (synaptic) source activity. Our dipolar sources were oriented in the posterior to anterior direction, aiming to capture this synaptic source activity.

### 4.2 Phase locking between the brain, spinal cord, and muscle during tonic contraction

Using the spinal cord region of interest identified by the source analysis, we then evaluated phase coupling between the brain, spinal cord, and muscle. We observed significant coupling between the brain and muscle, and between the brain and spinal cord, but not between the spinal cord and muscle. Coupling patterns were largely similar across participants and for right and left contraction.

For contraction of the right hand, coupling between brain and muscle activity was present in the beta band and was evident in all three participants. This is consistent with work showing that sensorimotor cortical and muscle activity are coupled at beta frequencies during maintained contraction of distal muscles using the coherence metric [14], [16], [17]. This cortico-muscular coherence has been strongly associated with descending activity in the corticospinal pathway [42], [43], and sensory input is known to have a modulatory effect [44], [45]. This cortico-muscular functional connectivity indicates that rhythmic activity is transmitted through the spinal cord, where it can impact and be impacted by spinal neurons [20]. This provides a point of departure for our further analyses; importantly we are now able to put these findings into context with concurrent measurements from the spinal cord. Note however that although we observed significant cortico-muscular coupling across participants for the right hand contraction, it did not reach statistical significance for left hand contraction (discussed in 4.4).

Brain and spinal cord activity also exhibited phase locking that was was significant and replicable across the frequency range of interest. Phase locking was generally evident both at lower frequencies (< 10 Hz) and at beta frequencies. One possible, albeit speculative, interpretation for these results is that the beta band phase coupling in particular, as a metric similar to coherence, reflects activity in the corticospinal pathway, at least in part [18], [42], [43].

Interestingly, phase locking between the spinal cord and muscle was negligible across participants for both right and left contraction. We might interpret this in the context of previous primate studies [19], [20] investigating coherence between spinal local field potentials and muscle activity. This work found that the prevalence of spinal LFP-EMG pairs showing significant coherence was relatively low (∼20 % of LFP-EMG pairs across two monkeys) [20], which may be linked to lack of phase coupling we observed.

One potential interpretation of this absence of phase coupling between the spinal cord and muscle, despite its presence between the other nodes of the network, is that transmission throughout this network is not a simple, passive process, and that the mode of communication may change between the different nodes. Another contributing factor may be that we are detecting activity in distinct populations of neurons in the different analyses. Future work using generative modelling and directed/effective functional connectivity analysis could certainly help clarify the characteristics of the underlying system but for now we speculate that it may be a result of the integrative nature of spinal cord processing.

### 4.3 Amplitude envelope coupling between the brain, spinal cord, and muscle

Beyond phase-locked interactions, we also found amplitude envelope interactions between all of the nodes in our network that were largely reproducible within and across participants.

The within-frequency amplitude envelope coupling we observed was generally focused on frequencies from 5-10 Hz, suggesting coupling between the slow dynamics of rhythmic activity in the brain, spinal cord, and muscle. During weak, maintained contraction – as is the case in our task – small movement discontinuities give rise to low frequency fluctuations in peripheral signals, including the EMG envelope. It has been demonstrated that this type of slow fluctuation in peripheral signals is coupled to sensorimotor cortical activity, termed ‘Cortico-kinematic coupling’ [46], [47], [48] and is linked to proprioceptive feedback [49] resulting from these changes in muscle force and length. Consequently, one interpretation of our envelope coupling is that it is capturing the dynamics of proprioceptive processing throughout the sensorimotor network. Future work to investigate the direction of coupling in this system may help to elucidate these claims.

Finally, we also detected significant coupling of amplitude envelopes across frequencies. The frequencies contributing to this coupling varied between participants and signal pairs; however, the most prominent and consistent features were generally peaks below 15 Hz. The presence of cross-frequency coupling in our data is in agreement with increasing evidence supporting cross-frequency coupling in the sensorimotor system, both in ascending and descending pathways, during tasks similar to our tonic thumb contraction task [50], [51], [52], although the precise characteristics of this type of functional connectivity are not yet well described. Cross-frequency coupling in the sensorimotor system has been associated with transmission in multi-synaptic pathways [52], [53]. Within-frequency coupling may dominate in monosynaptic pathways, like the direct corticospinal projections, whereas increasing synaptic layers lead to less linearity and more cross-frequency coupling [53], [54]. Consequently, the presence of cross-frequency amplitude envelope coupling throughout the network may be a signature of transmission and integration in pathways with multiple synaptic layers.

In sum, the key contribution of this paper is the concurrent imaging of activity in the brain and spinal cord at a millisecond time scale. We detected significant interactions between brain and spinal cord activity, and between spinal cord and muscle activity, in agreement with the known physiology of sensorimotor pathways. The neural coupling between these structures yielded heterogeneous functional patterns (i.e. modes of coupling) with different spectral signatures. This may hint at the underlying transfer properties of each node, and signify distinct modes of large-scale neural transmission in the sensorimotor system. Taken together, we suggest that the complex functional connectivity patterns are a hallmark of the spinal cord’s role in integration.

### 4.4 Methodological considerations

Critical to collection and analysis of OPM data is noise reduction [55]. We took several steps to reduce interference in the data: during the experiment, participants lay very still, only making small thumb movements, minimizing interference from movement through the background field and vibration of the MSR. In our analysis, temporal filtering reduced high and low frequency interference and line noise, synthetic gradiometry reduced environmental interference, and spatial filtering was used to remove heartbeat artefacts. The source reconstruction analysis also provided a spatial filtering function, further removing unwanted environmental and physiological noise. We additionally used reference sensor signals as a null space in our functional connectivity analysis for added noise immunity.

We modelled current flow in the spinal cord as a grid of current dipoles in an infinite homogenous volume. This model has the benefit that it is relatively robust to co-registration errors between anatomy and sensors because of its simplicity. More accurate and complex forward models, e.g. boundary element models of the torso, combined with more precise co-registration to individual spinal anatomy, will help to improve resolution and accuracy in future work. A noteworthy challenge in this context will be accurately modelling the detailed anatomical structures in the torso and their movement due to the heartbeat and breathing.

It is worth noting that although the majority of functional connectivity results were similar for right and left thumb contraction, the cortico-muscular phase coupling we observed only reached statistical significance in the beta band for the right hand contraction, but not for the left hand. One possible explanation for this is that precision was generally lower for left compared to right hand contraction (see Supplementary Table 2). Also, asymmetry in cortico-cortical coupling during motor tasks has been previously noted and suggested to relate to the left hemispheric dominance in right handers [56]. All other results were however largely reproducible within (right and left contraction) and between participants.

OPM based systems give us the flexibility to study the spatiotemporal dynamics of the nervous system as a whole, which is virtually unprecedented. Interactions between different regions of the CNS and the periphery are at the core of voluntary movement control, and OPM-based systems uniquely allow us to study this functional integration directly and non-invasively. In this work, we kept participants still to minimize interference, but when we have a better understanding of the signals we can record from the spinal cord, as well as the possible sources of interference, we aim to record activity during larger-scale, natural movements. This is already possible for brain activity recordings [55] and will represent an important step towards naturalistic and global imaging of the sensorimotor system.

## 5. Conclusion

In this work, we show that OPMs can be used to non-invasively image spinal cord current flow during voluntary hand contraction, and that endogenous, rhythmic spinal cord activity is significantly coupled with brain and muscle activity. Modes of communication (in terms of phase and/or amplitude envelope coupling) between these structures were heterogenous, suggesting differences in the neural populations and content that is propagated. This demonstrates the utility of OPMs for high-fidelity, spatiotemporal imaging of the entire central nervous system, which has the potential to fundamentally advance our understanding of how the nervous system communicates in health and disease.

## Data availability statement

The datasets generated and analysed during the current study will be made available upon reasonable request to the corresponding author.

## Author contribution statement

M.E.S, S.B., and G.R.B. devised the project. G.C.O, R.C.T, T.W., S.M., T.T., N.A., R.S., M.E.S., and G.R.B. wrote software. All authors contributed to the interpretation of the results and to the writing and editing of the manuscript.

