## Supplementary material for "Non-invasive evidence for rhythmic interactions between the human brain, spinal cord, and muscle"

#### 1. Sensor distribution

| Participant | Gen-2 neck | Gen-3 neck | Gen-2 head | Gen-3 head |
|---|---|---|---|---|
| A | 6 | 18 | 13 | 0 |
| B | 12 | 18 | 12 | 1 |
| C | 14 | 19 | 12 | 2 |

**Supplementary Table 1**: Number of Gen-2 (dual axis) and Gen-3 (triaxial) sensors in each participant's neck and head cast.

#### 2. Task performance

| Participant | Right | Left |
|---|---|---|
| A | 7.5± 3.2 | 7.3 ± 2.9 |
| B | 6.6 ± 3.3 | 9.1 ± 4.4 |
| C | 6.4 ± 3.2 | 10.6 ± 6.2 |

**Supplementary Table 2**: Task performance (precision) during tonic contraction. Values are mean and standard deviation for the root mean square error of the produced rectified, smoothed EMG vs. target level (AU).

### 3. Within-frequency phase locking and amplitude envelope coupling for left contraction

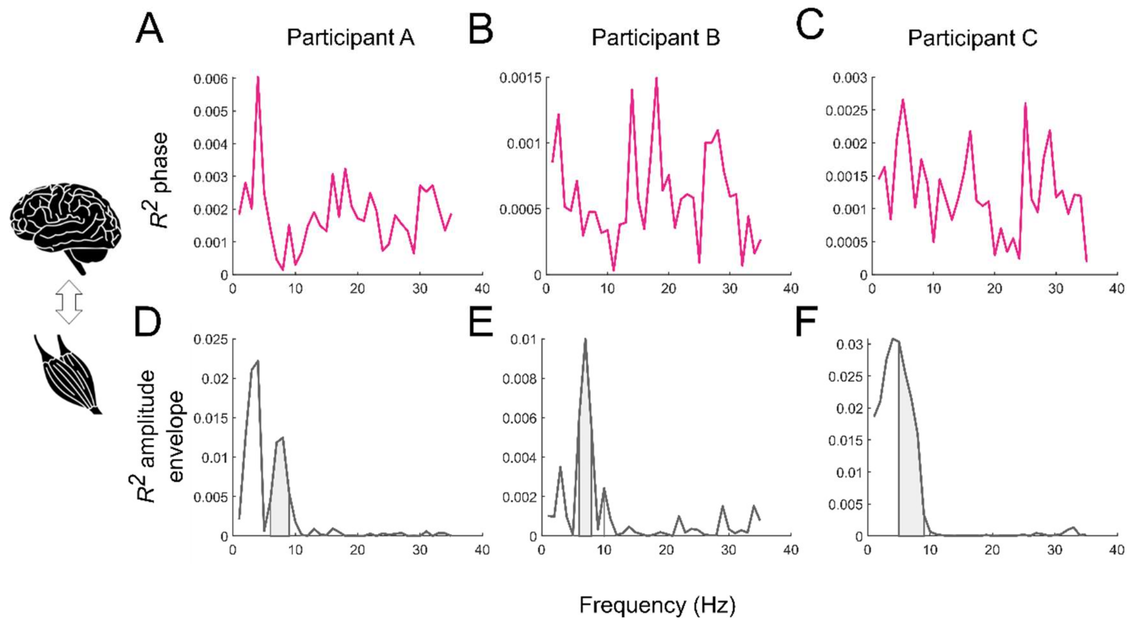


**Supplementary Figure 1: Functional connectivity between brain and muscle activity during left thumb contraction.** $R^2$ for within-frequency phase locking (**A-C**). Vertical lines and shaded areas indicate FDR corrected $p<0.05$. **D-F**: $R^2$ for within-frequency amplitude envelope coupling. Y-axes are scaled differently to clearly show spectral features.

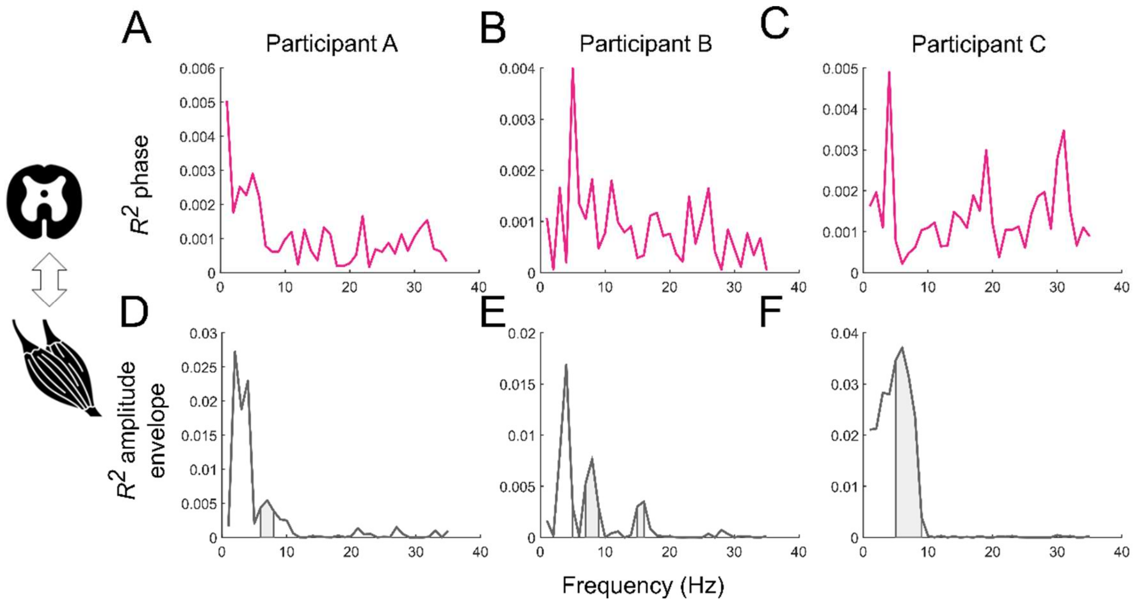


**Supplementary Figure 2: Functional connectivity between spinal cord and muscle activity during left thumb contraction.** $R^2$ for within-frequency phase locking (**A-C**). Vertical lines and shaded areas indicate FDR corrected $p<0.05$. **D-F**: $R^2$ for within-frequency amplitude envelope coupling. Y-axes are scaled differently to clearly show spectral features.

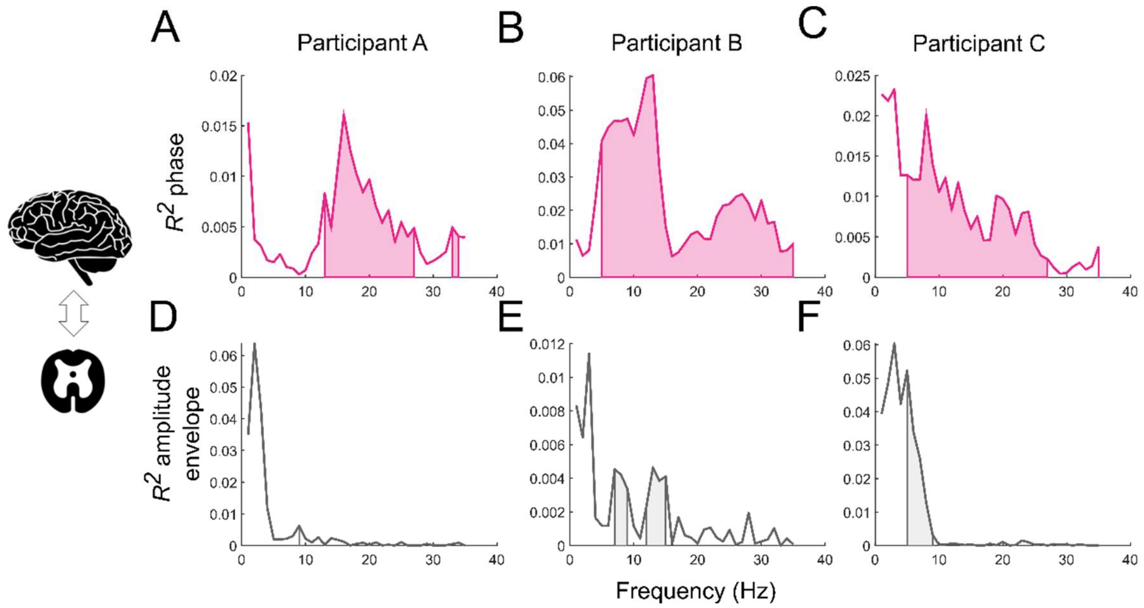


**Supplementary Figure 3: Functional connectivity between spinal cord and brain activity during left thumb contraction**. $R^2$ for within-frequency phase locking (**A-C**). Vertical lines and shaded areas indicate FDR corrected $p<0.05$. **D-F**: $R^2$ for within-frequency amplitude envelope coupling. Y-axes are scaled differently to clearly show spectral features.

### 4. Cross-frequency envelope coupling and canonical vectors

The statistical significance of the cross-frequency envelope coupling during right contraction is reported in the main text. Here, we report corresponding results for left contraction. For all participants and all signal pairs, we found a significant cross-frequency amplitude envelope coupling ($p < 0.001$) in our frequency range of 5-35 Hz during contraction of the left hand.

Below, we visualize the coupling results for both right and left contraction as the normalized canonical vectors for each signal pair. The normalization was performed according to (Haufe et al., 2014) eq. 6, where the weights/filters in the backward model are the canonical vectors and the latent factors are the canonical variates.

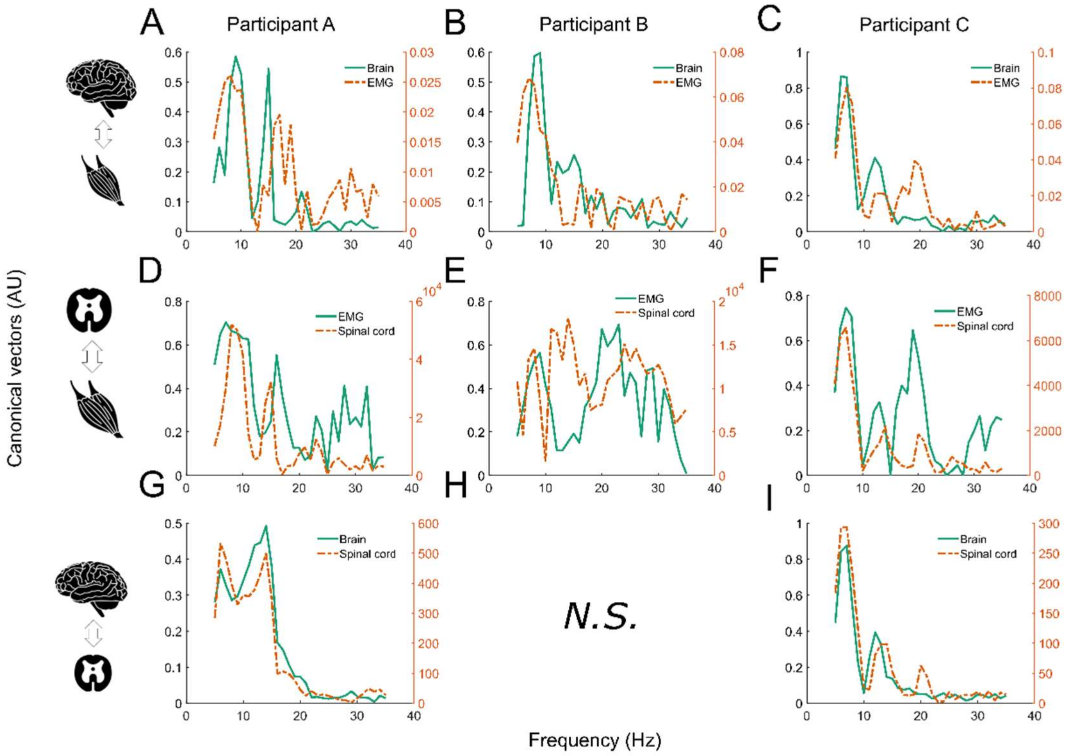


**Supplementary Figure 4: Canonical vectors for cross-frequency envelope coupling during right contraction.** The canonical vectors are shown for the first significant mode of coupling for each participant and signal pair. Values reflect the contribution from each frequency to the coupling between the two signals. Two second axes are used because the vectors are on different scales. The scales of the vectors do not have neurophysiological significance. *N.S* indicates that the coupling was not significant.

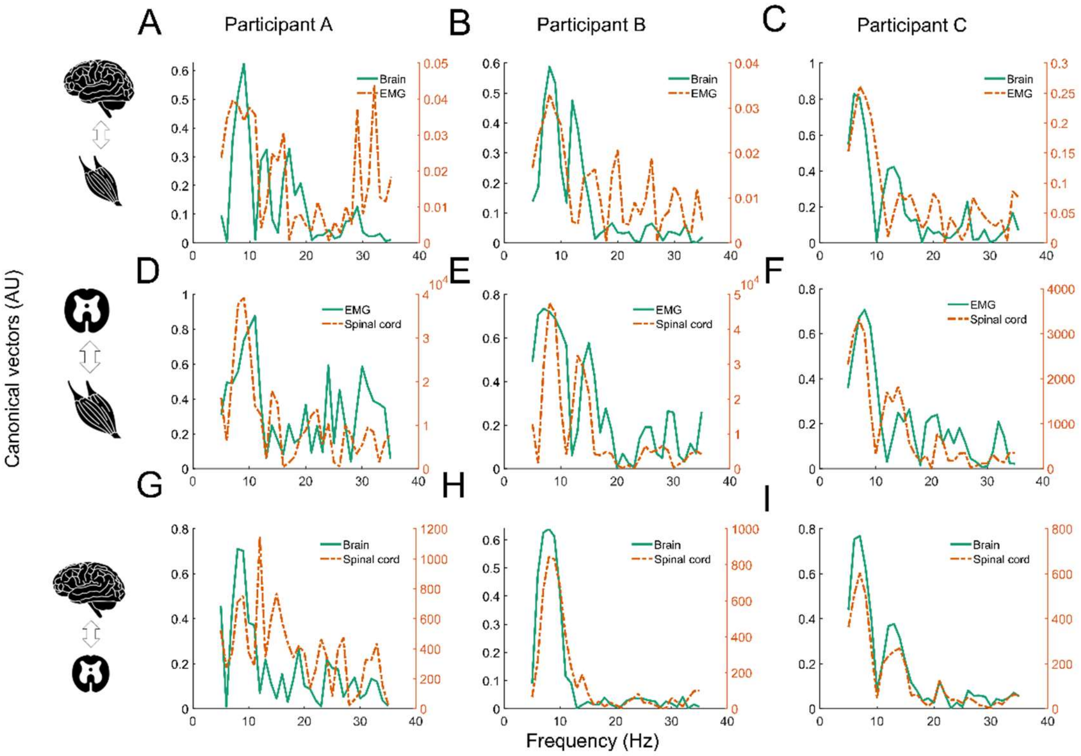


**Supplementary Figure 5: Canonical vectors for cross-frequency envelope coupling during left contraction.** The canonical vectors are shown for the first significant mode of coupling for each participant and signal pair. Values reflect the contribution from each frequency to the coupling between the two signals. Two second axes are used because the vectors are on different scales. The scales of the vectors do not have neurophysiological significance.

### 5. Comparison of coherence with CVA-based phase locking

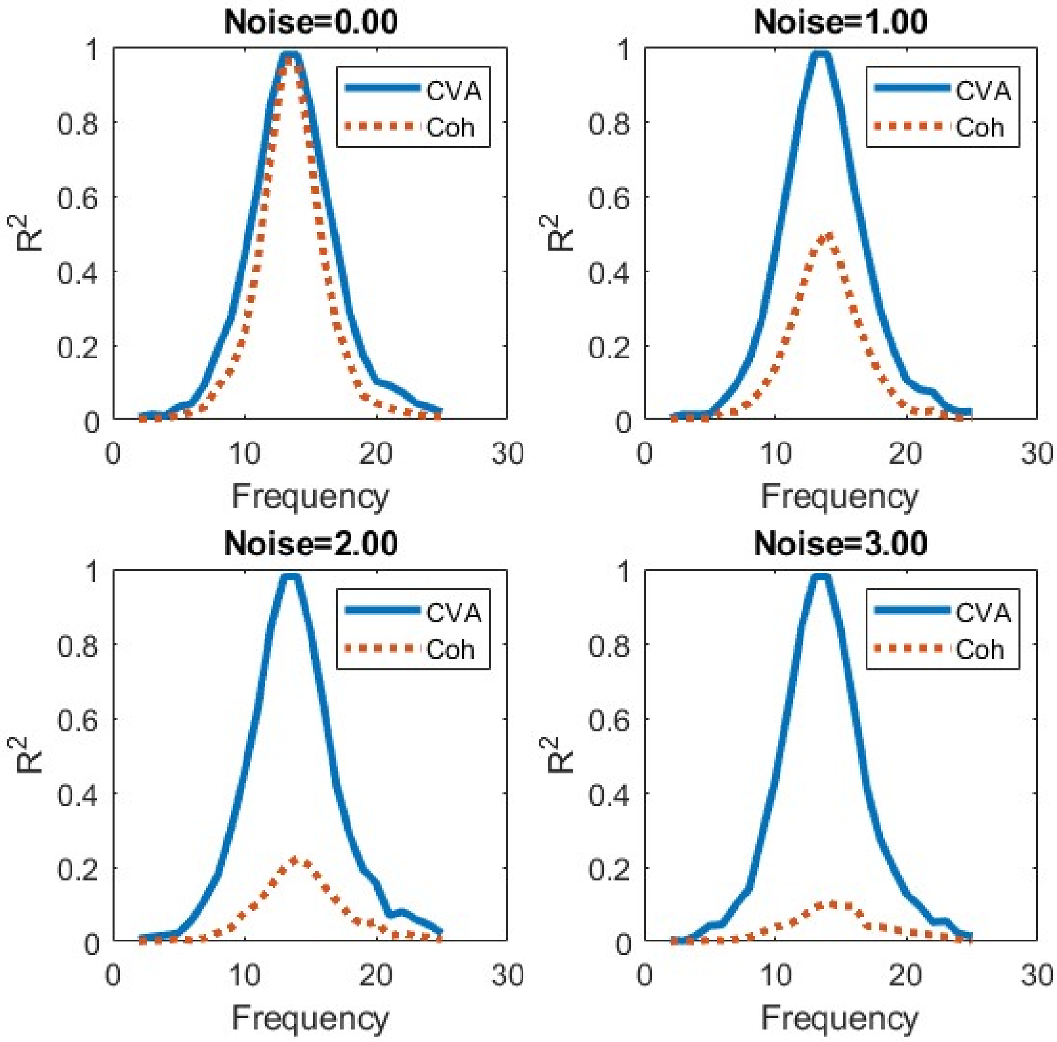


**Supplementary Figure 6. Comparison of CVA-based phase locking and coherence.** To compare the performance of CVA to the traditional coherence metric, we simulated two signals; *x* and *y*. The imaginary/cosine component of *y* was constructed to share a direct phase relationship with the real/sine component at a specific frequency. Signal *y* also contained an 'interferer', i.e., an additional real/sine component at the same frequency not directly related to *x*. In panels above, the amplitude of this interferer (*Noise*) is varied between 0 and 3.0. When there is no interferer (*Noise*=0) the coherence (dotted) and CVA (solid) estimates accord. However, as the amplitude of the interferer is increased, coherence decreases (as the effective phase at this frequency is being perturbed) whereas CVA (which deals with real and imaginary components independently) remains constant.

Sample MATLAB code (phase_cva_example.m) to create this figure is available at https://github.com/meaghanspedden/CVA_coh_sim.git
